# Predicting Toxicity and Bioactivity of the Chemical Exposome: A Case Study for the Blood Exposome Database

**DOI:** 10.1101/2025.11.10.687755

**Authors:** Ankita Dutta, Dinesh Barupal

## Abstract

Humans are exposed to thousands of chemicals throughout their life. Many of these chemicals are detected in blood and have been catalogued in the Blood Exposome Database. Comprehensive hazard assessment of a chemical requires time-consuming and costly lab experiments using animal or cell-lines, which cannot be easily scaled up to the chemical exposome, highlighting the urgent need for computational approaches that can prioritize chemicals based on toxicological information. In this study, we trained direct message passing neural networks (D-MPNN) models using the Chemprop framework chemical structure and bioactivity data from 9,458 compounds profiled in the U.S. EPA’s Tox21 program across 148 quantitative high-throughput screening assays. Additionally, we trained a complementary model using chemical structures (n=264,601) labeled with known UN GHS classifications for acute oral toxicity. Both models demonstrated strong predictive performance, with average AUCs exceeding 0.80 for 47 Tox21 assays. We applied these 48 models to 58,673 chemicals from the Blood Exposome Database to predict bioactivity and the GHS hazard classification, enabling scalable in-silico prioritization of understudied chemical exposures for further toxicological investigations. Data and code are available at https://zenodo.org/records/17560382 and https://github.com/idslme/exposome-toxicity-prediction.

## Introduction

The exposome includes all the non-hereditary agents^1,2^ that can adversely affect molecular processes, including gene regulations, epigenetic processes, signaling pathways, and metabolic pathways, which are involved in the onset and progression of the human diseases that have a minimum genetic predisposition^1^ such as cancer, diabetes, and Alzheimer’s disease. Studying Exposome is high-priority research by governmental and private organizations to improve human health^3,4^. The toxic potential of an exposome agent is a critical attribute for nominating them^5^ for more legal and safety scrutiny and reviews. The chemical exposome includes all the chemicals with a toxicity^6^, for example carcinogens, neurotoxins or endocrine disruptors^7^. These chemicals can have both natural and synthetic origins. The toxic substance control act (TSCA) of the USA^8^ has listed over 80,000 chemicals, both natural and synthetic, that are utilized in the consumer industry. Many of these chemicals have been detected in human biospecimens, suggesting that humans are exposed to them and there is a need to understand the toxicity of these chemicals. Human biospecimen, particularly blood, serve as reservoirs of circulating chemical exposures and are therefore widely studied in toxicological hazard identification^9,10^. The Blood Exposome Database ^11^ contains over 50,000 chemicals that have been reported for blood specimens. This database is continuously growing as more chemicals are detected for blood specimens using untargeted and targeted assays developed using highly sensitive mass spectrometry instruments. This resource provides a critical reference framework to improve the annotation and interpretation of compounds identified through MS based analysis. However, a majority of these chemicals lack toxicity or bioactivity data.

Predictive toxicology includes development and application of machine learning models^12^ that can predict toxicity and bioactivity endpoints for a chemical^13^. With the availability of curated and high-quality toxicity and bioactivity datasets^14^, machine learning frameworks such as Chemprop^15^, and better computer hardware including graphical processing units, it is feasible to develop and apply predictive toxicology models for a large list of chemicals such as the Blood Exposome Database. Since generating experimental data for each chemical’s toxicity is expensive, laborious and resource-demanding for large chemical lists, AI/ML models for predicting toxicity^16^ are practical tools to add toxicity indications for chemicals. Furthermore, such models can aid in prioritization^16^ of chemicals that require further experimentation and scrutiny under regulatory policies, such as the TSCA.

In this work, we apply the Directed Message Passing Neural Network (D-MPNN) architecture, implemented in the Chemprop framework^15^, to develop predictive models for toxicological risk assessment. Here, we report the development of 47 models to predict bioactivity endpoints and an additional model to predict the acute toxicity of a chemical. We have utilized publicly available datasets including Tox21, UN-GHS classification^17^, Blood Exposome Database, and the Chemprop framework to train these models. The work prioritized several chemicals from the Blood Exposome Database that may be toxic to the human body and require further testing using laboratory studies, including NAMs. The goal of this work is to enable high-accuracy prediction of biological endpoints given the chemical structure of compounds within the human exposome.

## Methods

Our workflow combines bioactivity, toxicity and chemical exposome databases, and uses state-of-the-art machine learning models to predict toxicity and bioactivity classification for a chemical structure (Figure 1).

**Figure 1.**
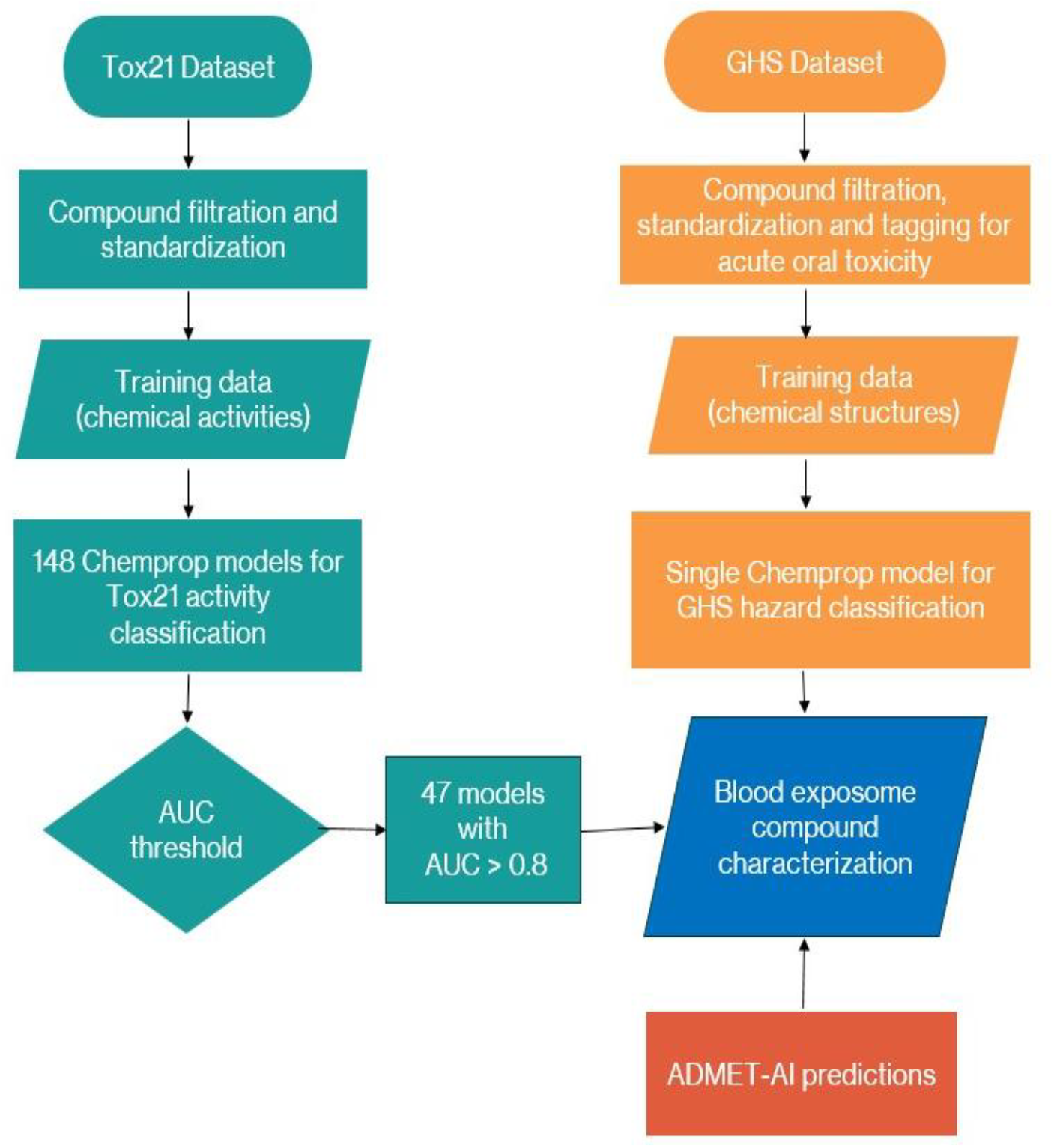
Overview of the chemical datasets and machine learning models developed and utilized in this study.

### Training Data preparation

Tox21 (tox21.gov) is a comprehensive and publicly available bioactivity dataset generated by screening approximately 8,500 compounds across more than 70 bioassays. These assays were prioritized by a consortium of US Government agencies to develop methods and resources for chemical safety and toxicity analyses. The dataset has been widely used for predictive modeling of chemical toxicity through structure–activity relationship analysis^14^. Two separate datasets were obtained in October of 2024 from the NCBI PubChem database: (1) A summary of all bioassays included in the Tox21 study (https://pubchem.ncbi.nlm.nih.gov/source/824) was downloaded in .csv format referred to as the “Tox21 data” hereafter (2) A “bioactivities.tsv” (https://ftp.ncbi.nlm.nih.gov/pubchem/Bioassay/Extras/bioactivities.tsv.gz) dataset that contained the activity outcomes across all bioassays that are available in the PubChem database. This dataset consisted of 13 columns which can be grouped as follows : (i) Assays and compound type identifiers (AID, SID, SID group and CID), (ii) Bioassay outcome based on compound activity (Activity outcome, Activity Name, Activity Qualifier and Activity Value), (iii) Publication related to the bioassay and biological targets (Protein Accession, Gene ID target, Tax ID and PMID).

To prepare the training dataset, we first obtained the list of bioassays included in the Tox21 dataset. The bioactivities.tsv dataset was then subsetted for compound CIDs and bioactivities that were included in the Assay ID (AID) that were part of the Tox21 study. The data set was filtered by removing any datapoint with incomplete information such as missing Compound IDs (CIDs). Datapoints with “Inconclusive” or “NaN” in the *Activity Outcome* column were also discarded. The resulting dataset was filtered to only include bioassays with known target proteins and only bioassays with assigned Uniprot or NCBI protein ascension numbers were considered for model training. SMILES for the compounds are obtained from PubChem identifier exchange service and added to the dataset by matching unique CIDs. Any datapoint without a valid SMILES was also discarded. The final dataset consisted of 148 total bioassays and 9852 unique CIDs that met the above filtration criteria.

Prior to model training, the dataset was split into 148 subsets corresponding to each bioassay. Experimental activity outcomes from the Tox21 bioassays were encoded as binary labels (active = 1, inactive = 0). Across all 148 subsets, the SMILES strings were standardized and desalted using RDKit libraries. Finally, redundant data with multiple activity values corresponding to a single compound SMILES were removed. These redundant activity values corresponded to varying concentrations of the same compound being tested within a bioassay. Since the goal of this work is to classify compounds irrespective of their concentrations, in this work, we only retain a compound SMILES when the compound is found to be *active* in an assay of interest. Only the first occurrence of a compound was retained when the compound is found to be either active or inactive across all occurrences.

### GHS Oral Toxicity Classification

The Globally Harmonized System of Classification and Labelling of Chemicals (GHS) is a United Nations-developed international standard for classifying chemical hazards and communicating information on labels and Safety Data Sheets. It is a globally agreed upon system that harmonizes chemical hazard classification and labeling, ensuring consistent and clear communication of chemical risks across countries and industries. The GHS classification system recognizes three key hazard classes for chemicals – Environmental, Health and Physical hazards^17^. As of October 2024, a total of 264,641 compounds were catalogued in the PubChem database with a GHS classification. In this study we focus on compounds with an acute oral toxicity warning under the GHS classification. To prepare the training data we obtained an exhaustive list of compounds with any GHS classification in addition to a list of compounds with only an acute oral toxicity warning. The list of compounds and their associated chemical properties were obtained in an .sdf file format directly from the PubChem website (https://pubchem.ncbi.nlm.nih.gov/classification/#hid=83). Internal codes were then utilized to produce a unified list of compounds where each compound was given a binary tag [0, if the compound does not have an acute oral toxicity warning but is included in the GHS compound list; 1, if the compound is present in the GHS compound list and has an acute oral toxicity warning]. Of the 264,641 compounds 40 metal ion SMILES were incompatible with RDkit due to unresolved electronic charges^18^. Our final dataset processed for model training consisted of 264,601 total compounds of which 154,602 had an acute oral toxicity tag.

### Blood Exposome Database

The Blood Exposome Database (www.bloodexposome.org) was compiled in previous work (https://zenodo.org/records/8146024) and is utilized here to exclusively test the efficacy of the DMPNN models for compound toxicity classification. Similar to the Tox21 and GHS datasets, the compound SMILES within the Blood Exposome Database were standardized and desalted. Redundant smiles were discarded and only the first occurrence of redundant SMILES was retained for further analysis. The final Blood Exposome Database consisted of 52,055 compound SMILES.

### Chemprop model training

Chemprop is a deep learning framework that leverages graph neural networks (GNNs) to predict molecular properties directly from SMILES representations. By modeling molecules as graphs, Chemprop captures atom-level interactions through message passing, enabling accurate classification without relying on hand-crafted descriptors^15^. Compared to traditional machine learning models, molecule graphs are shown to perform better for predictive toxicology^19-21^. A total of 148 individual molecular activity classification models were trained using the Chemprop framework, each corresponding to one of the 148 bioassays included in the Tox21 dataset. Additionally, a complementary model was trained to predict Globally Harmonized System (GHS) classification outcomes.

Prior to model training, the compound lists associated with each of the 148 individual assays in the Tox21 dataset, as well as the list used for GHS classification were randomly partitioned into training (75%) and validation (25%) subsets. Molecular structures were provided as canonical SMILES strings, which were tokenized and transformed into graph-based representations using Chemprop’s default featurization—where atoms are represented as nodes and bonds as edges. These molecular graphs served as input features for the DMPNN architecture implemented within Chemprop. Chemprop training and predictions were performed using a single GPU node. Further hardware and software specifications are provided in subsequent subsection. Model performance was assessed on the held-out validation sets using standard classification metrics, with particular emphasis on the area under the receiver operating characteristic curve (ROC-AUC).

### ADMET-AI Predictions

ADMET-AI is an open-sourced machine learning platform. The source code for which was downloaded directly from the github page (https://github.com/swansonk14/admet_ai). ADMET-AI has been widely used to predict ADMET (Absorption, Distribution, Metabolism, Excretion, and Toxicity) properties from molecular string representations (e.g., SMILES). It is built on the Chemprop-RDKit architecture, which extends the original Chemprop framework by integrating additional molecular descriptors and features computed using RDKit, a cheminformatics toolkit. This enhancement allows the model to leverage both learned molecular graph representations and expert-crafted features for improved prediction accuracy^13^. The ADMET-AI models were trained on datasets from the Therapeutics Data Commons (https://tdcommons.ai/). ADMET-AI prediction was used to supplement our findings and gain further understanding of the toxicological characteristics of the compounds within the Blood Exposome Database. The Clintox predicted by the ADMET-AI were selected to show the relevant toxicity endpoints.

### Model performance metrics

The predictive performance of the classification models was evaluated by calculating the area under the receiver operating characteristic curve (AUC-ROC). The AUC-ROC was calculated by integrating the ROC curve, which plots the true positive rate (TPR), TPR = TP / (TP + FN), against the false positive rate (FPR), FPR = FP / (FP + TN) across all possible probability thresholds. In practice, the AUC was computed using the trapezoidal rule implemented in *scikit-learn’s* “roc_auc_score” function, which provides a threshold-independent measure of model discrimination between active and inactive compounds. The optimal classification threshold was determined using Youden’s J statistic, defined as, J = sensitivity + specificity – 1, were, Sensitivity (true positive rate) = TP / (TP + FN) and Specificity (true negative rate) = TN / (TN + FP). The threshold corresponding to the maximum J value along the ROC curve was selected, and predictions were binarized at this cutoff to compute accuracy and additional performance metrics.

### Morgan fingerprinting and PCA plotting

Molecular representations were generated using Morgan circular fingerprints with a radius of 2 and a bit vector length of 1024, as implemented in RDKit. Each compound’s SMILES string was converted to a binary fingerprint encoding the presence or absence of atom-centered substructures within the defined radius. Principal component analysis (PCA) was then performed on the fingerprint matrix using scikit-learn’s implementation (sklearn.decomposition.PCA). The dimensionality of the 1024-bit fingerprints was reduced to two principal components, which were then used for visualization of compound clustering.

### Software and computing specifications

All computations were performed on a workstation running Ubuntu Linux 22.04 (kernel 5.15.0-138-generic) with an Intel(R) Xeon(R) Gold 6342 CPU @ 2.80 GHz, 503 GB RAM, and an NVIDIA A100 GPU (40 GB memory). Models were implemented in Python 3.11.11 using Chemprop v2.1.0, RDKit v2024.9.4, scikit-learn v1.6, and PyTorch Lightning v2.5.0.post0.

All the generated datasets are available at Zenodo (https://zenodo.org/records/17560382).

## Results

### Bioactivity model selection

We started our analysis by finding which bioactivity models had acceptable accuracy. We trained 148 bioactivity prediction models using the Tox 21 dataset. The distribution of active versus inactive compounds across all 148 bioassays included in the final Tox21 dataset (Figure 2) was not uniform. The number of compounds tested in these bioassays ranged between 5000 to 8000 unique compound CIDs. The number of active CIDs showed had a standard deviation of 481.42 across the 148 bioassays. Assays with less than 3,000 CIDs tested produced a lower AUC and worse predictive performance. A similar pattern was also observed when the ratio of active to inactive CIDs was highly skewed (Figure 2). Of the 148 classification models, 47 models met the AUC>0.80 criterion and were deemed acceptable and used to classify the compounds in the Blood Exposome Database. Of the 52,055 unique CIDs in the Blood Exposome Database, 4,188 CIDs were also tested in the Tox21 program. False positive rates for this subset of compounds were relatively low across all 47 high accuracy classification models (Figure 3).

**Figure 2.**
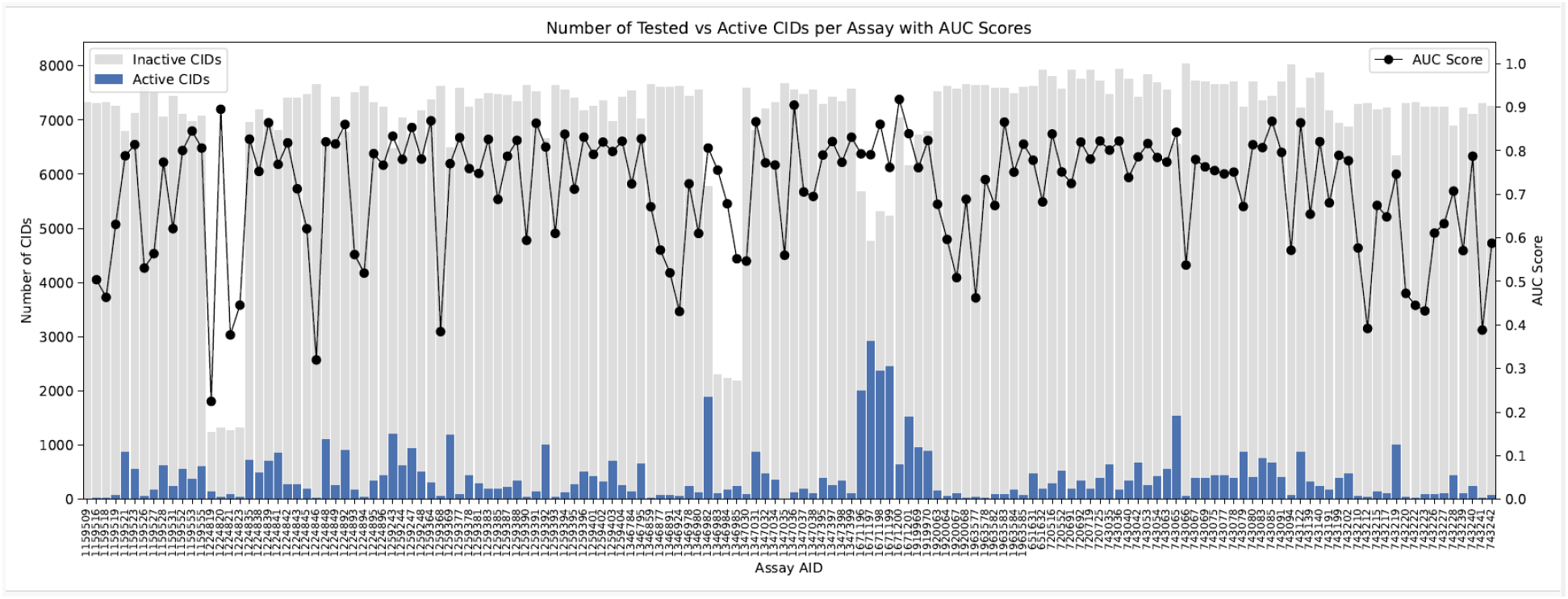
Distribution of the number of unique Compound IDs across 148 Tox21 bioassays, with active CIDs in blue and Chemprop model AUCs shown as black dots.

**Figure 3.**
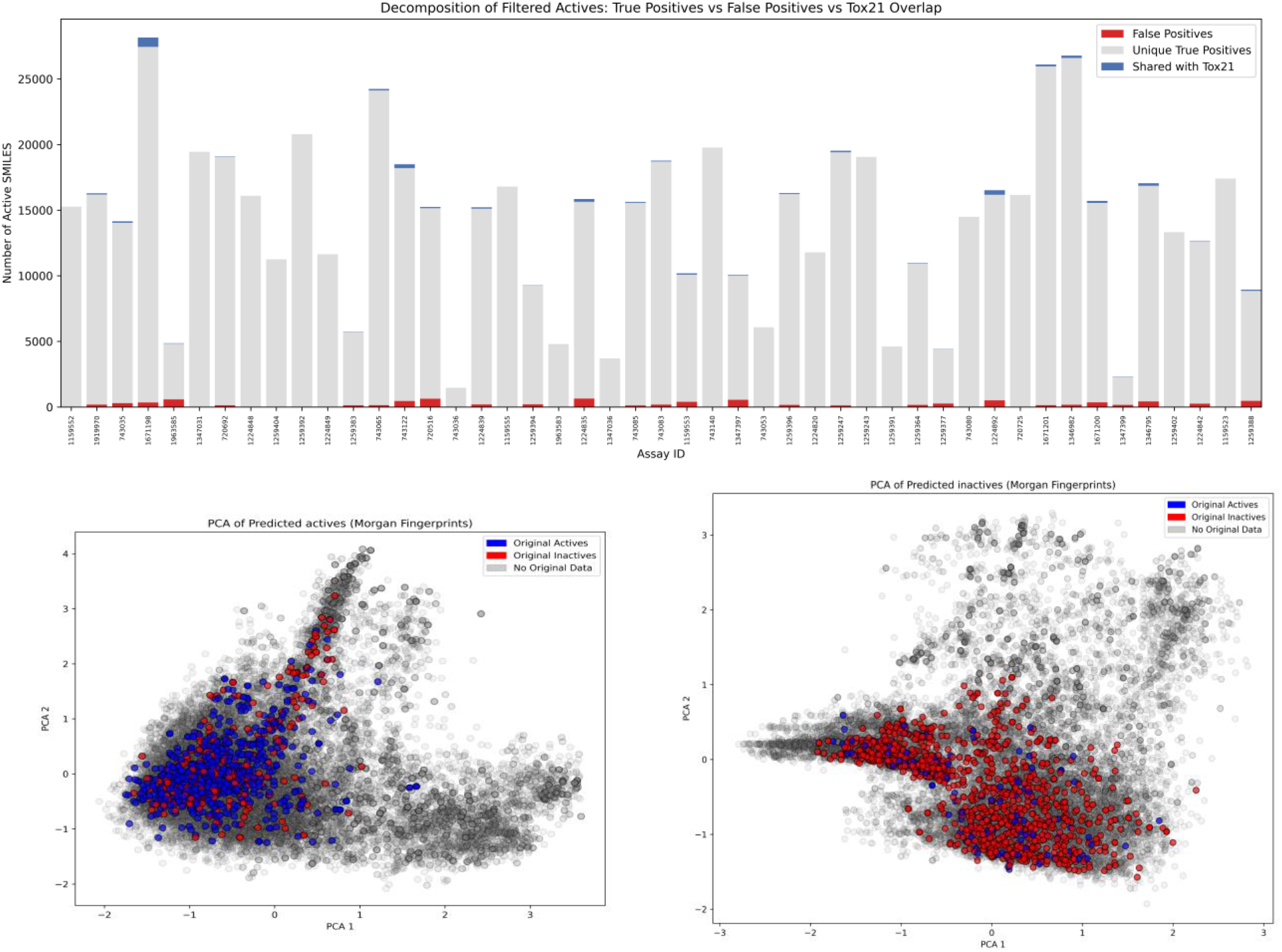
(Top panel) Stacked bar chart showing all predicted active CIDs (Grey bar), true positives (blue bar) and false positives (red bar) across 47 Tox21 assays. (Bottom left panel) PCA plot of all unique CIDs predicted to be active and (Bottom right panel) PCA plot of all unique CIDs predicted to be inactive in AID 1671198.

To investigate the underlying reasons for the misclassification, we performed Principal Component Analysis (PCA) on the Morgan fingerprints of the compounds (Figure 3) to check the domain compatibility. Specifically, we visualized the PCA projections of compounds categorized as true positives, false positives, and all predicted positives for AID 1671198 where the number of true positives observed is larger than false positives. The PCA plots revealed that true positives and predicted positives were clustered within the same region of the chemical space between Tox21 and the Blood Exposome Database, suggesting that Chemprop consistently identified a specific structural motif or fingerprint pattern. False positive and false negative are mainly focused on the chemical space region that had minimum overlap between the Tox21 and the Blood Exposome Database.

For compound to be prioritized as potentially toxic and be recommended for further experimental testing, the active classification of a compound should be supported by multiple bioactivity classification models. The frequency with which a compound is predicted to be active was tallied (Figure 4), to prioritize the probably toxic compounds. 2,5’-Bithieno[3,2-b]furan (PubChem CID 71619088) is predicted to be active across all 47 classification models (Figure 4). We also note that a large portion of the compounds in the Blood Exposome Database are classified as active in only 1 to 6 bioassays. Fused aromatic ring structural motif was a common feature observed among the top seven CIDs most frequently predicted to be active (Figure 4).

**Figure 4.**
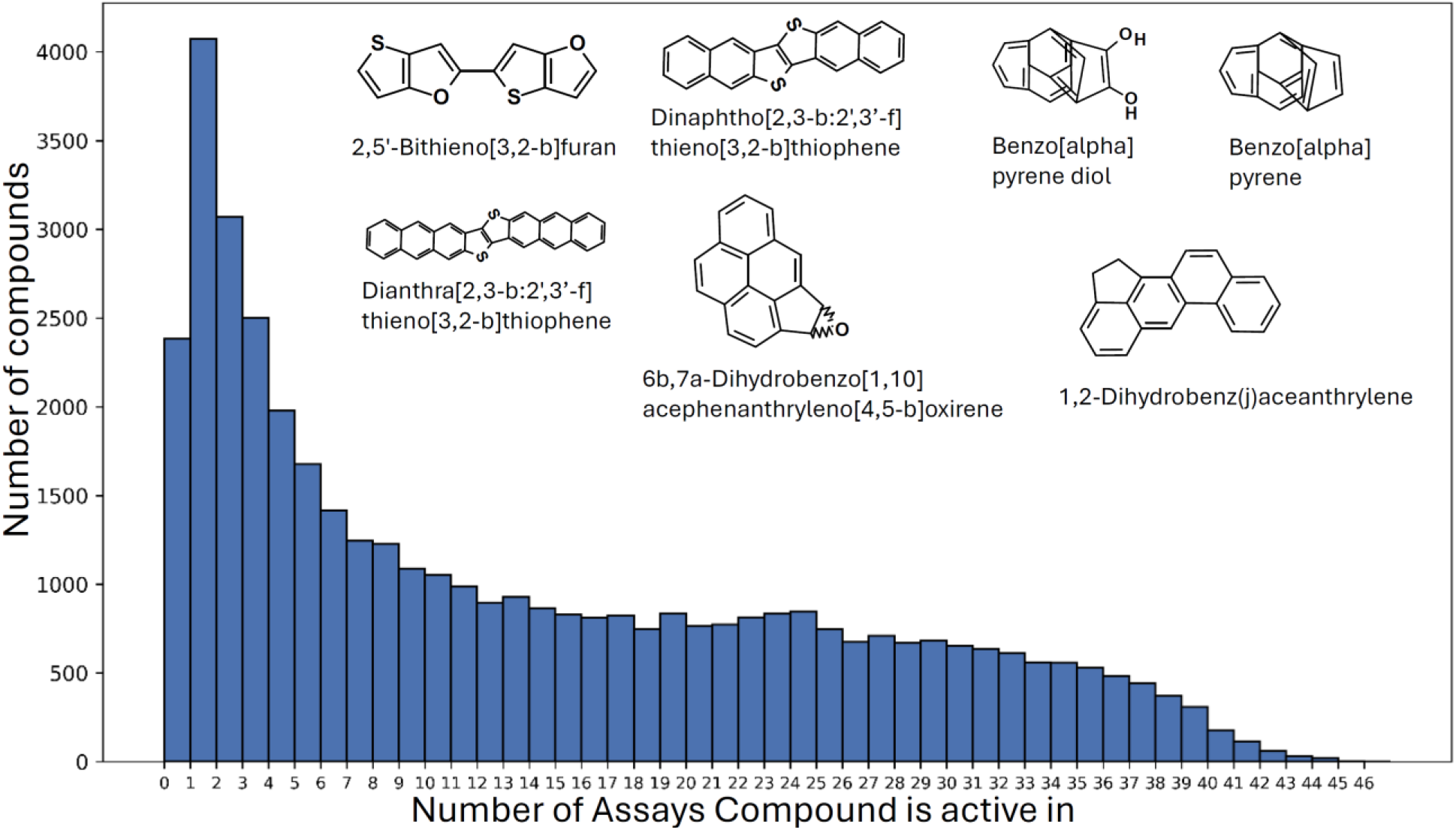
Histogram shows the frequency with which unique CIDs from the Blood Exposome Database are classified as active across 47 Tox21 bioassay models. Chemical structures of the top seven compounds from the Blood Exposome Database are predicted to be active across the widest range of Tox21 bioassay models.

Next, we tested whether the DMPNN-based models can predict GHS hazard classifications. We prepared a complementary GHS classification model. The model showed high accuracy with an AUC of 0.81 (Figure 5). Subsequently, the model was applied to predict the GHS classifications of all compounds in the Blood Exposome Database. Overlapping chemical space, which included the compounds that were present in both the GHS and Blood Exposome Database, was used to check the accuracy of the model predictions (Figure 5). We noted that the frequency of accurate predictions (true positives and true negatives) is higher than inaccurate predictions (false positive and false negative) for the overlapping compounds. 2,5’-Bithieno[3,2-b]furan is also classified as orally toxic by the GHS model, which supported our previous findings and prioritized this compound for experimental testing.

**Figure 5.**
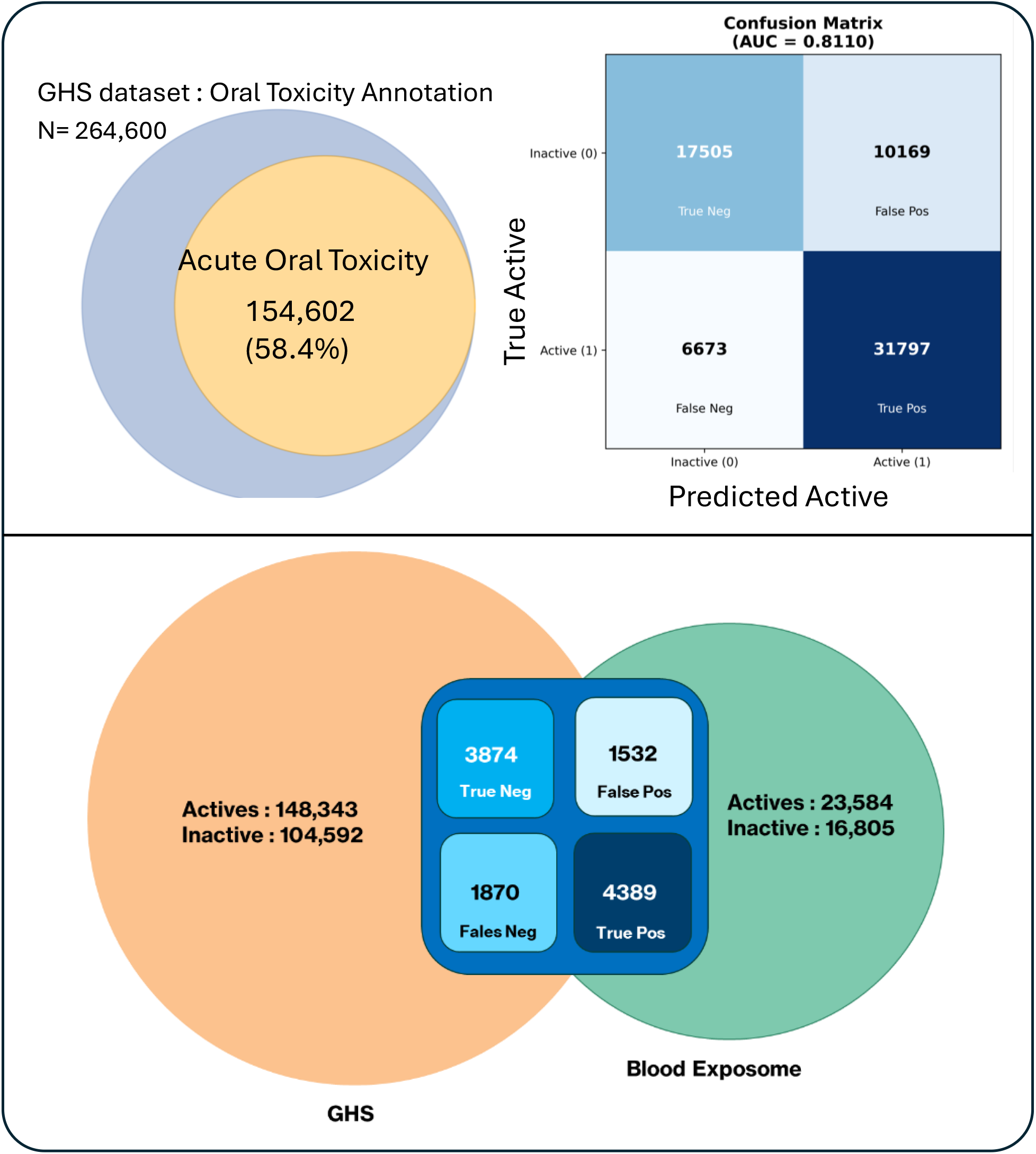
(Top left panel) Venn diagram showing the total number of unique CIDs with known GHS hazard classification (blue circle) and the subset of unique CIDs with an acute oral toxicity label (yellow circle). (top right panel) Confusion matrix of the compound classification prediction of CIDs in the holdout validation set. (bottom panel) Venn diagram showing the overlap between the CID in the Blood exposome and GHS datasets. Overlapping compounds are broken down in a confusion matrix format to show the accuracy of classification predictions.

Next, we predicted ADMET endpoints for the compounds in the Blood Exposome Database. 3,678 unique CIDs from the Blood Exposome Database were covered in both the Tox21 and GHS databases. For these compounds, ClinTox predictions were generated using the open-source models available through the ADMET-AI package (Figure 6).

**Figure 6.**
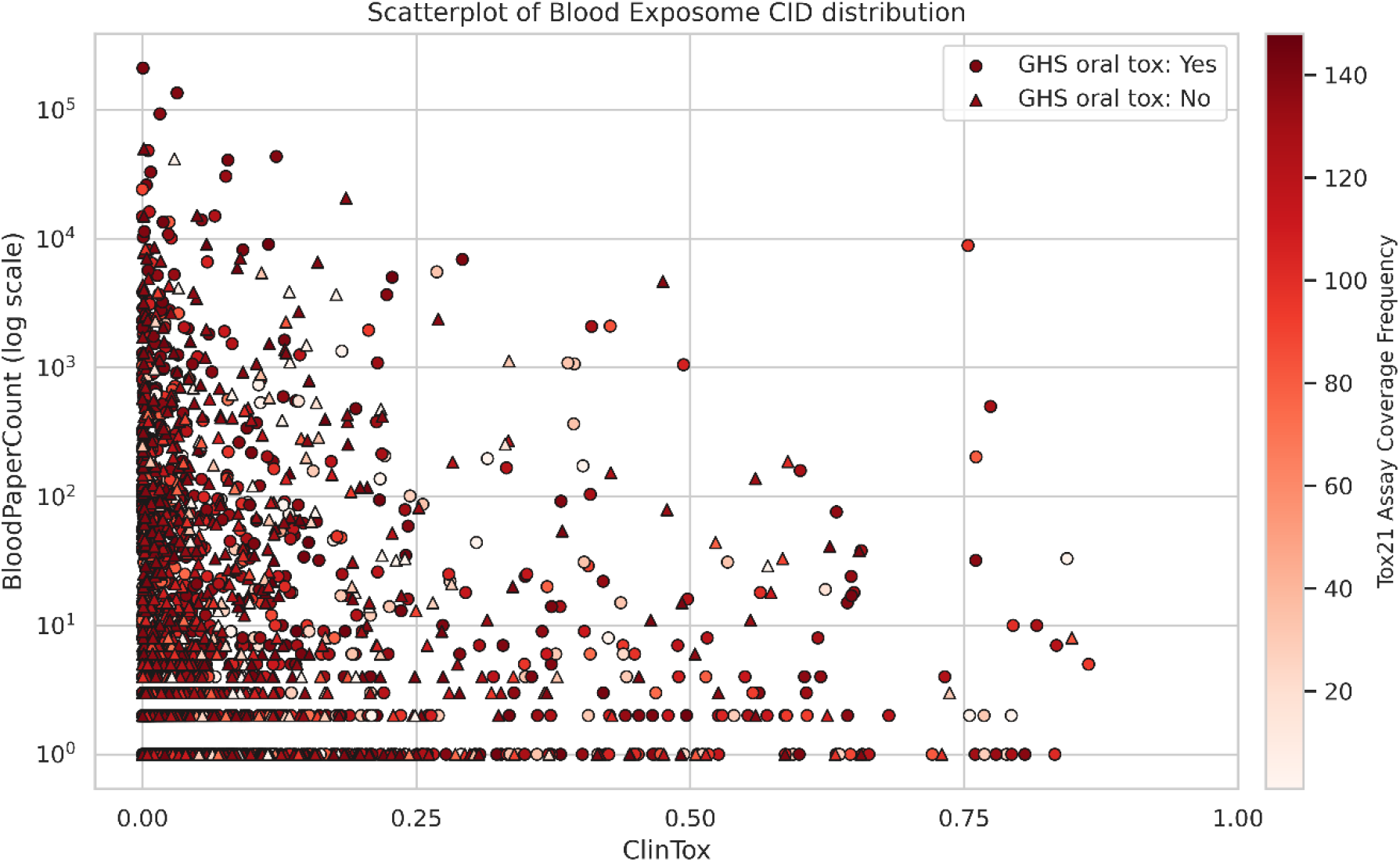
Scatterplot of unique CIDs in the Blood Exposome Database that are also covered in both the Tox21 and GHS datasets. ClinTox (Top panel) values are probability predicted using external ADMET-AI models. A probability of 1.0 indicate the highest possible toxicity. Compounds with an acute oral toxicity classification in the GHS dataset are shown in blue circles and all other compounds are shown in red triangles. The color gradient of each datapoint represents the frequency with which the compound is covered across the 148 Tox21 bioassays.

Visualization of ClinTox metrics in relation to publication frequency revealed that many overlapping compounds cluster toward lower ClinTox values. In contrast, compounds with higher ClinTox scores are frequently classified under the acute oral toxicity category by the GHS system, yet their testing frequency within the Tox21 program remains relatively low.

We applied the previously trained Chemprop models for 47 bioactivity, GHS Oral Toxicity, along with ADMET-AI predictions to enrich the toxicity data for all Blood Exposome Database compounds. It revealed a consistent trend in which most compounds cluster toward lower ClinTox values (Figure 7). Compounds with higher ClinTox scores were frequently predicted to have an acute oral toxicity tag according to the GHS classification model. Notably, we observed a cluster of compounds with ClinTox scores between 0.50 and 0.75 that are well represented in published literature. These compounds are often predicted to carry an acute oral toxicity classification under the GHS system and exhibit a higher frequency of being predicted as active in the Tox21 models.

**Figure 7.**
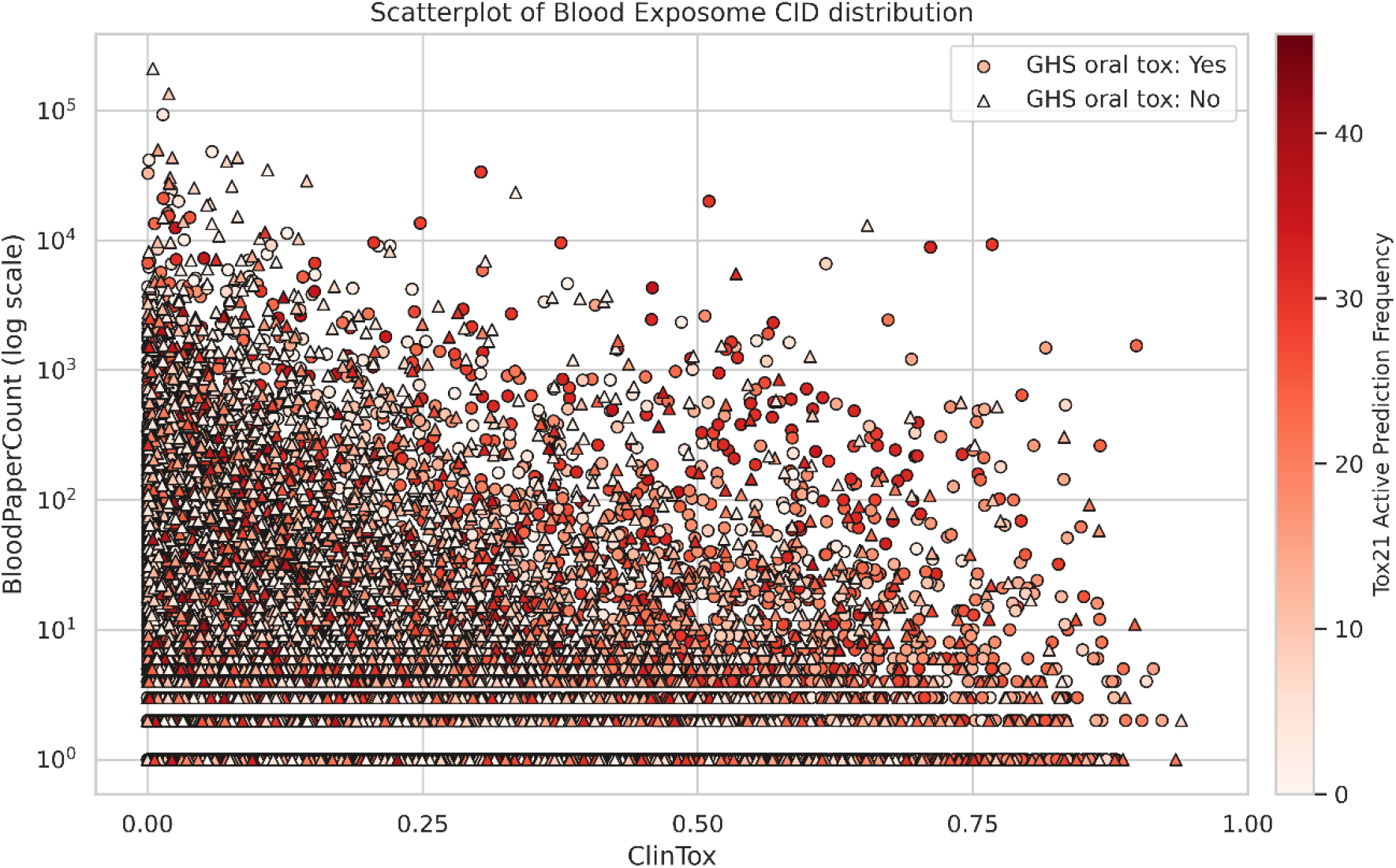
Scatterplot of unique CIDs in the Blood Exposome Database that lack coverage in either the Tox21 or GHS datasets. ClinTox (Top panel) and DILI (bottom panel) values are probability predicted using external ADMET-AI models. A probability of 1.0 indicated the highest possible toxicity. Compounds predicted to have an acute oral toxicity classification in the GHS dataset are shown in blue circles and all other compounds are shown in red triangles. The color gradient of each datapoint represents the frequency with which the compound predicted to be active by the 47 Tox21 bioassays models with AUCs > 0.80.

To further support these predictions, we checked the top-ranked compounds for published literature on toxicity. Many of these compounds are natural products and were tested anticancer drugs which are known to cause cell death. For example, oxoglaucine had known hepatotoxicity^22^, Blebbistatin and Wogonin are generally cytotoxic^23,24^, whereas chemicals with benzofurans and thiopene groups are known to cause cellular toxicity^25,26^. These examples support that our integrated modelling approach can be used for prioritizing toxic chemicals in the Blood Exposome Database for further experimental validations. For this top 100 chemicals that have very limited literature data can be given a higher priority.

Together, our results highlight the utility of machine learning–based methods in expanding the toxicological information for an important biomedical knowledgebase, the Blood Exposome Database. By integrating existing experimental and curated data within the Chemprop framework, we identified previously underrepresented compounds that exhibit signatures of potential toxicity. ClinTox predictions further supported our observations. Overall, this work demonstrates how predictive modeling can complement limited assay data, enhance toxicological profiling, and guide future experimental prioritization within large-scale exposomic datasets.

## Discussion

The chemical exposome^27^ is vast and evolving as new natural and synthetic compounds are generated every year. It is widely accepted that there is a need to develop computational predictive methods^16,28-31^ that can flag chemicals that might be harmful^32^ to human health. In this report, we demonstrate the use of machine learning models to predict bioactivity and toxicity of the compounds in the Blood Exposome Database^11^. DMPNN-based Chemprop models^15^ can extend coverage to compounds that remain untested, offering a scalable and systematic approach for exposome wide toxicity profiling. This will be helpful in prioritizing compounds in the Blood Exposome Database that need further attention for biomonitoring, experimental toxicology studies and additional in-silico predictions.

Only about 7% of chemicals from the Blood Exposome Database have existing regulatory data through programs such as the U.S. EPA Tox21 and the UN-GHS. To address this gap, we developed a machine learning-based framework to enrich toxicological information for compounds that remain untested, complementing the chemical prioritization efforts^6,33^. To this end, we developed 47 high-priority classification models, each corresponding to a known Tox21 bioassay, in addition to a complementary GHS acute oral toxicity classification model. Through our ensemble modeling strategy, we were able to strengthen the confidence of predictions made by any individual model. We highlighted a compound, 2,5’-Bithieno[3,2-b]furan, which exhibited the highest frequency of toxic classification across all models tested in this study. We recommend that top 100 compounds ranked by our strategy should be prioritized for further studies, but the enriched database can be used to flag other chemicals that are relevant for the consumer industry or environmental pollution. Taken together, our ensemble modeling framework enables the ranking of chemicals in the human blood exposome based on predicted toxicity classifications, thereby supporting compound prioritization for further experimental evaluation.

However, several limitations remain. Broader assay representation and a diverse distribution of bioactivities within datasets will be essential to improve model robustness and generalizability. Moreover, a deeper understanding of the molecular mechanisms underlying toxicity, such as protein-level interactions, is necessary to fully interpret computational predictions. This will be addressed in our future work, where we aim to investigate these mechanisms through ensemble docking studies against key metabolic proteins, such as cytochrome P450s^34^. Overall, this study highlights the importance of integrated multiple machine learning models that predict complementary information to evaluate the hazard potential of the chemical exposome.

## Funding

The work was in part supported by U24ES035386, R24ES036917, R01ES035478, P30ES023515, UL1TR004419, R01ES032831 and R01ES033688.

## Conflict of interest

The authors declare no competing financial interest.

## Author’s contribution

AD and DB planned the study, implemented the scripts, prepared the results and drafted the manuscript. All authors have reviewed the manuscript’s content.

## Data and code availability

Data and code are available at https://zenodo.org/records/17560382 and https://github.com/idslme/exposome-toxicity-prediction

## Notes

https://zenodo.org/records/17560382

### Competing Interest Statement

The authors have declared no competing interest.

